# METACLUSTER^*plus*^ - an R package for probabilistic inference and visualization of context-specific transcriptional regulation of biosynthetic gene clusters

**DOI:** 10.1101/2022.04.11.487835

**Authors:** Michael Banf

## Abstract

Fungi and plants reveal widespread occurrences of metabolic enzymes co-located on the chromosome, some already characterized as being biosynthetic pathways for specialized metabolites, such as terpenes synthesizing enzyme clusters in *Lotus japonicus* and *Arabidopsis thaliana*. These clusters display context-specific co-expression of clustered enzymes, indicating a shared transcriptional response in a spatial and condition specific manner, and co-regulation due to promoter binding by shared transcription factors may be one way to facilitate coordinated expression. To enhance our understanding of context-specific transcriptional gene cluster regulation, we redefine and augment this probabilistic framework, labelled METACLUSTER^*plus*^, integrating gene expression compendia, context-specific annotations, biosynthetic gene cluster definitions, as well as gene regulatory network architectures. Further, it provides a set of appealing and intuitive visualizations of inferred results for analysis and publication. METACLUSTER^*plus*^ is available at https://github.com/mbanf/MetaclusterPlus.

## 1. Introduction

Plants as well as microbial organims produce a variety of compounds denoted as specialized metabolites to cope with environmentsal challenges but the biosynthetic pathways for many of these compounds have not yet been elucidated [15]. Recent studies in plants [6, 13, 10, 14, 16] revealed a widespread occurrence of metabolic enzymes that collocate in the chromosome. This offers an intriguing possibility for uncovering new biosynthetic pathways encoded by these metabolic gene clusters. To this end, co-expression analysis can provide valuable insights as characterized specialized metabolic pathways and clusters exhibit high degrees of co-expression among their enzymes [13, 10, 17]. Moreover, the expression patterns of experimentally characterized gene clusters indicate spatial and condition specificity, such as enzymatic genes of clusters synthesizing terpenes in *A. thaliana* and *L. japonicus* [11, 7, 8, 17].

To facilitate a convenient and context-specific activity analysis of metabolic gene clusters, we recently proposed a probabilistic framework [1], denoted METACLUSTER, which automatically identifies conditions and tissues associated with inferred gene clusters within a given differential gene expression compendium. However, of equal importance, in particular with the emergence of large-scale transcription factor binding data such as [2], is the elucidation of metabolic gene cluster transcriptional regulation, since it has been argued that one way to facilitate such coordinated gene expression may be co-regulation due to promoter binding by shared transcription factors [3]. Hence, to enhance our understanding of context-specific transcriptional gene cluster regulation, we redefine and augment this probabilistic framework, hence denoting it METACLUSTER^*plus*^, integrating gene expression compendia, context-specific annotations, biosynthetic gene cluster definitions, as well as gene regulatory network architectures. Cluster regulation is then inferred based on a series of statistical analyses, integrated via Fisher’s method [12], including statistical significance scores of metabolic cluster activity in a specific context, metabolic cluster enzyme co-regulation by a transcription factor within that context, as well as optional cluster evidence scores, such as enrichment of signature enzymes per cluster (see figure 1). METACLUSTER^*plus*^ may be applied to any organism, gene cluster descriptions, and differential gene expression datasets, thereby providing a valuable complementary framework to augment gene cluster inference approaches, such as PlantClusterFinder [13], antiSMASH [5], plantiSMASH [10], and PhytoClust [14], with additional layers of automated high-resolution functionality and transcriptional regulation inference. Further, it provides a set of appealing and intuitive visualizations of inferred results for eludication and publication.

**Figure 1:**
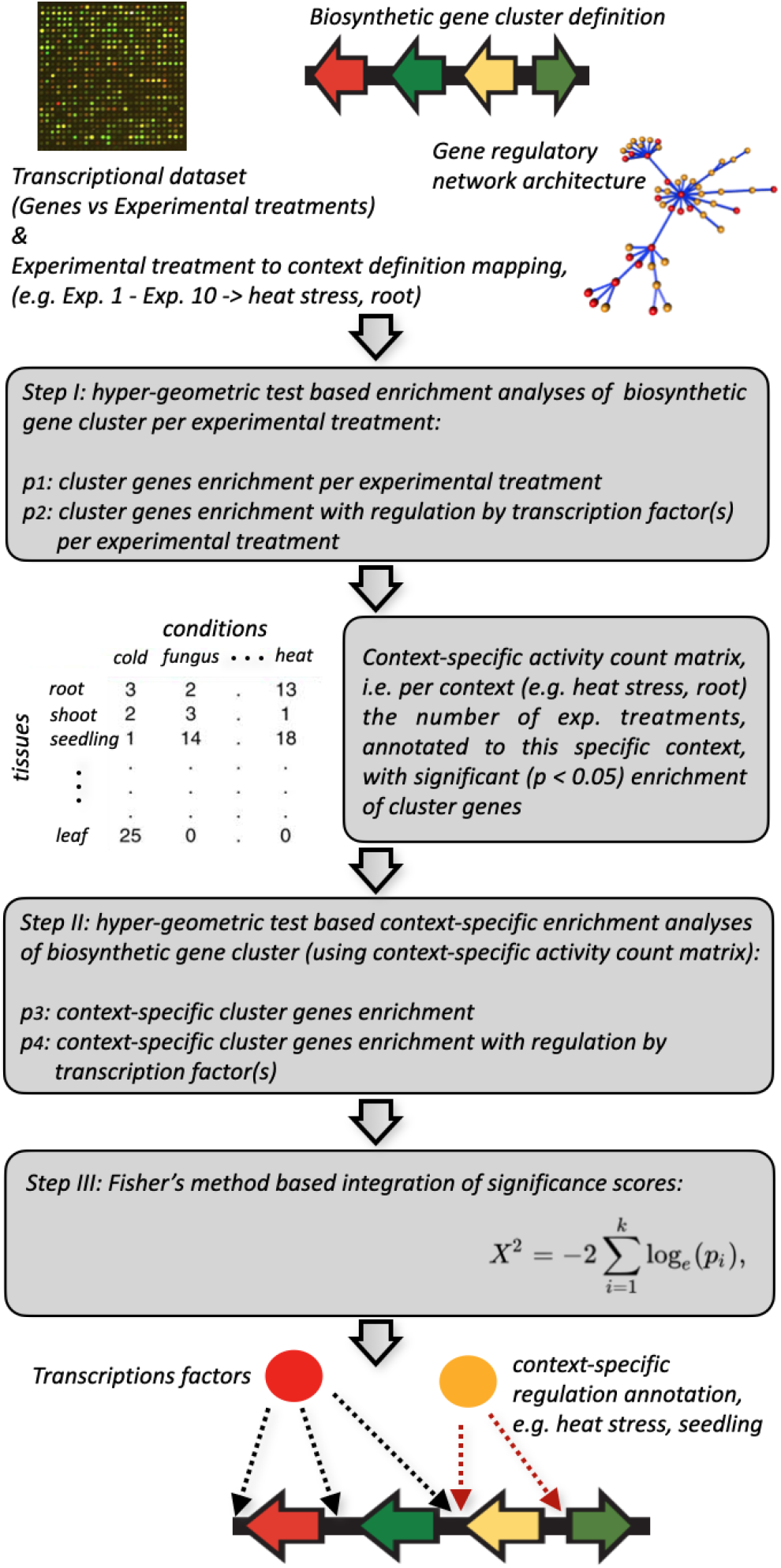
Overview of the probabilistic gen cluster regulation inference framework.

## 2. Methods

### 2.1. Inference of context-specific transcriptional activity and regulation

Schlapfer *et al*. [13] proposed a probabilistic ranking framework based on co-expression to identify sets of high confidence metabolic gene clusters and to prioritize clusters for experimental validation. This framework had been extended in our previous work [1] in order to allow for the identification of context-specific gene expression of metabolic gene clusters. For each gene cluster, a rank was introduced based on combining multiple evidence scores regarding coexpression among cluster genes and the cluster’s context specific transcriptional activity, all integrated using Fisher’s method [12]. While keeping the multiple evidence integration based approach, METACLUSTER^*plus*^ redefines the transcriptional activity inference in order to compensate for a potential weakness in the original framework. It further augments transcriptional activity analysis by another layer, that is the simultaneous inference of context specific transcriptional regulation.

Initially, as in [1], a differential gene expression dataset is constructed by retaining experimental treatments represented by gene expression profiles measuring gene expression responses of wild type to treatment and control conditions and computing the log of fold change difference between the mean of the treatment and control sample replicates. Two sample t-tests are performed per gene on each of the experiments to evaluate the significance of a gene’s differential expression between treatment and control, producing a ternary matrix *D* over all genes. For each gene and experimental treatment, an entry in *D* is assigned 1, -1 or 0 for statistically highly significant (*p* < 0.05) up-, down-, or non-significant differential expression.

Given *D*, our novel framework first estimates the probabilitiy *p*_*gc*_(*e*) of a gene cluster *gc* to be transcriptionally active in an experiment *e*. Since each experimental treatment *e* is annotated with a specific pair of condition *c* and tissue *t, e* may be defined as *e* := (*c, t*). We select experimental treatments in the differential expression matrix *D* that were statistically enriched in *gc* based on hyper-geometric test following a general hyper-geometric distribution 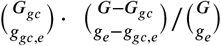 with *G* and *G*_*gc*_ denoting the total number of genes in the genome and the number of gene cluster genes, within experimental treatment *e. g*_*e*_ and *g*_*gc,e*_ represent the number of genes being differentially expressed in experimental treatment *e* and the subset of differentially expressed cluster genes in *e*, respectively. Following this hyper-geometric test, all experimental treatments with *p* ≤ 0.05 were selected. Using our manually curated mapping of experimental treatments to conditions and tissues, we then established a conditions vs tissue count matrix *C*_*gc*_ for further downstream context analysis (see figure 1). The context count matrix uses the association of individual experimental treatments *e* to the conditions *c* and tissues *t*, i.e. a pair of condition and tissue is incremented, if the corresponding treatment *e* for the cluster is significantly expressed (*p* ≤ 0.05). We then performed hyper-geometric tests to identify enrichment of *gc* for a specific context, i.e. a pair of (*c, t*), defined as probability *p*_gc_ (*c, t*) following the distribution 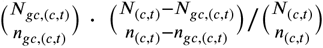 with *N*_(*c,t*)_ and *N*_*gc*(*c,t*)_ denoting the total number of all context pairs (*c,t*) and the number of all context pairs for a gene cluster *gc*. Accordingly, *n*_(*c,t*)_ and *n*_*gc*,(*c,t*)_ denote the number of a specific context pairs as well as the number of that context pair (*c, t*) for a specific gene cluster *gc*. Defining context specific activity in this manner, we also compensate for potential weakness in the original framework [1] where a rather artificial disentanglement of condition and tissue annotations was introduced due to the sequential approach that separately analyzed enrichment of conditions only first, thereby ignoring putatively conflicting tissues, and subsequently trying to add tissues. Further, as a beneficial side-effect, this further simplifies the whole process of transcriptional activity analysis and allows for a more unambiguous identification of specific cluster genes being as being transcriptional active compared to [1].

Next, we estimate *p*_*gc*_(*c, t,r*), which represents the probability of cluster *gc* to be transcriptionally active in a specific context (*c, t*) with a putative regulator for all regulators active in (*c, t*) with a minimum of two putative target genes of gene cluster *gc* being analyzed. Enrichment analysis is, again, based on a hyper-geometric distribution 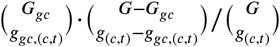 with *G* and *G*_*gc*_, with *t*_*r*,(*c,t*)_ and *t*_*gc,r*,(*c,t*)_ representing the number of targets of regulator in general and the number of targets genes expressed in experimental treatment (*c, t*), respectively. Next, we define context count matrix *C*_*gc,r*_ per gene cluster *gc* and putative regulator *r*. Again associated individual gene clusters and putative regulation to condition and tissue labels., we performed hyper-geometric test 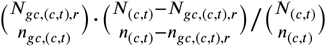, with *N*_(*c,t*)_ and *N*_*gc*,(*c,t*),*r*_ denoting the total number of contexts (condition and tissue) pairs and the total number context pairs for a gene cluster and associated regulator *r*, and a specific context (*c, t*). Accordingly, *n*_(*c,t*)_ and *n*_*gc*,(*c,t*),*r*_ denote the number of a specific context as well as the number of that context (*c, t*) for a specific gene cluster *gc* and regulator *r*.

Our method further allows for the integration of additional evidences, here for example the enrichment *p*_*gc*_(*sig*) of signature enzymes per cluster [13]. Integration of all individual evidence probability scores per gene cluster *gc* follows our previously proposed approach in [1] based on Fisher’s method [12] to estimate a combined p-value *p*_*gc*_(*r* ∈ (*c, t*)) to define a final score of involvement of regulator with gene cluster *gc*, given some condition *c* and tissue *t*.

### 2.2. Visualization of context-specific transcriptional activity and regulation

Aside a textual representation of the inferred gene cluster regulation (as illustrated in table 1), we equip our framework with chord graph based visualization per gene cluster that allows for i) transcription factor families vs conditions and treatments on a gene cluster level (see figures 2 and 4), as well as ii) transcription factor families vs conditions and treatments vs individual cluster genes on a cluster specific gene level (see figures 3 and 5). Numbers represent an actual count of associations between members of a transcription factor family and the gene cluster or cluster genes, respectively, given a specific condition and treatment.

**Table 1.** Selected example results of transcriptional activity and regulation inference for the thalianol cluster

| Transcription factor (family) | Treatment | Tissue | Regulated Genes |
| --- | --- | --- | --- |
| AT3G22760 (Trihelix) | salt | leaf | AT5G47950, AT5G48000, AT5G48010 |
| AT4G00730 (C2C2-Dof) | salt | root | AT5G47970, AT5G47990, AT5G48000, AT5G48010 |
| AT3G61150 (WRKY) | phosphate deprivation | shoot | AT5G47980, AT5G47990, AT5G48000, AT5G48010 |
| AT1G76890 (LBD) | phosphate deprivation | shoot | AT5G47980, AT5G47990, AT5G48000, AT5G48010 |
| AT4G14770 (NAC) | phosphate deprivation | shoot | AT5G47950, AT5G48000, AT5G48010 |
| AT5G47370 (NAC) | phosphate deprivation | shoot | AT5G48000, AT5G48010 |
| AT2G22430 (AP2/ERF-ERF) | phosphate deprivation | shoot | AT5G47980, AT5G47990, AT5G48000, AT5G48010 |
| AT2G30590 (C2C2-Dof) | pathogen | seedling | AT5G47950, AT5G47980 |
| AT3G01970 (WRKY) | pathogen | seedling | AT5G47950, AT5G47980, AT5G47990, AT5G48000 |
| AT5G47370 (MADS-MIKC) | pathogen | seedling | AT5G47950, AT5G47980, AT5G47990, AT5G48000 |
| AT3G21890 (HB-HD-ZIP) | auxin | root | AT5G47950, AT5G47980, AT5G47990, AT5G48000<br>AT5G48010 |

**Figure 2:**
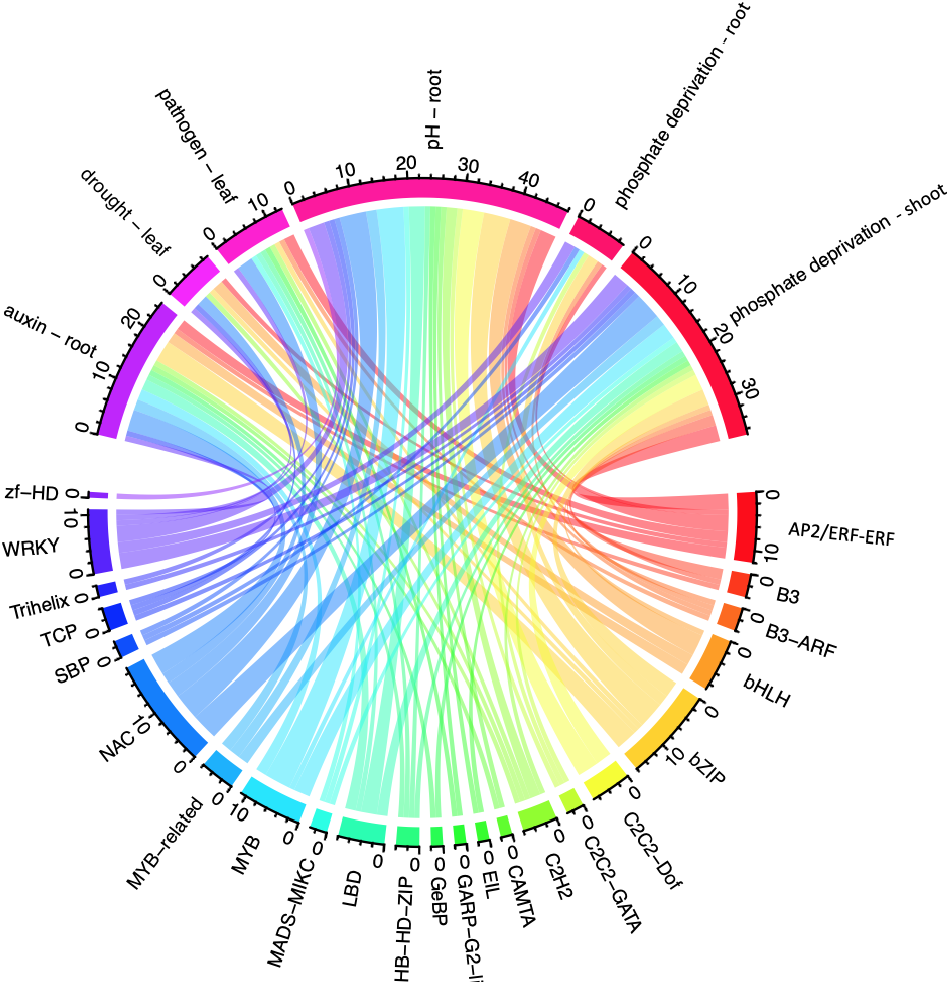
Gene cluster level visualization, i.e. transcription factor families vs conditions and treatments, of transcriptional activity and regulation of the marneral gene cluster.

**Figure 3:**
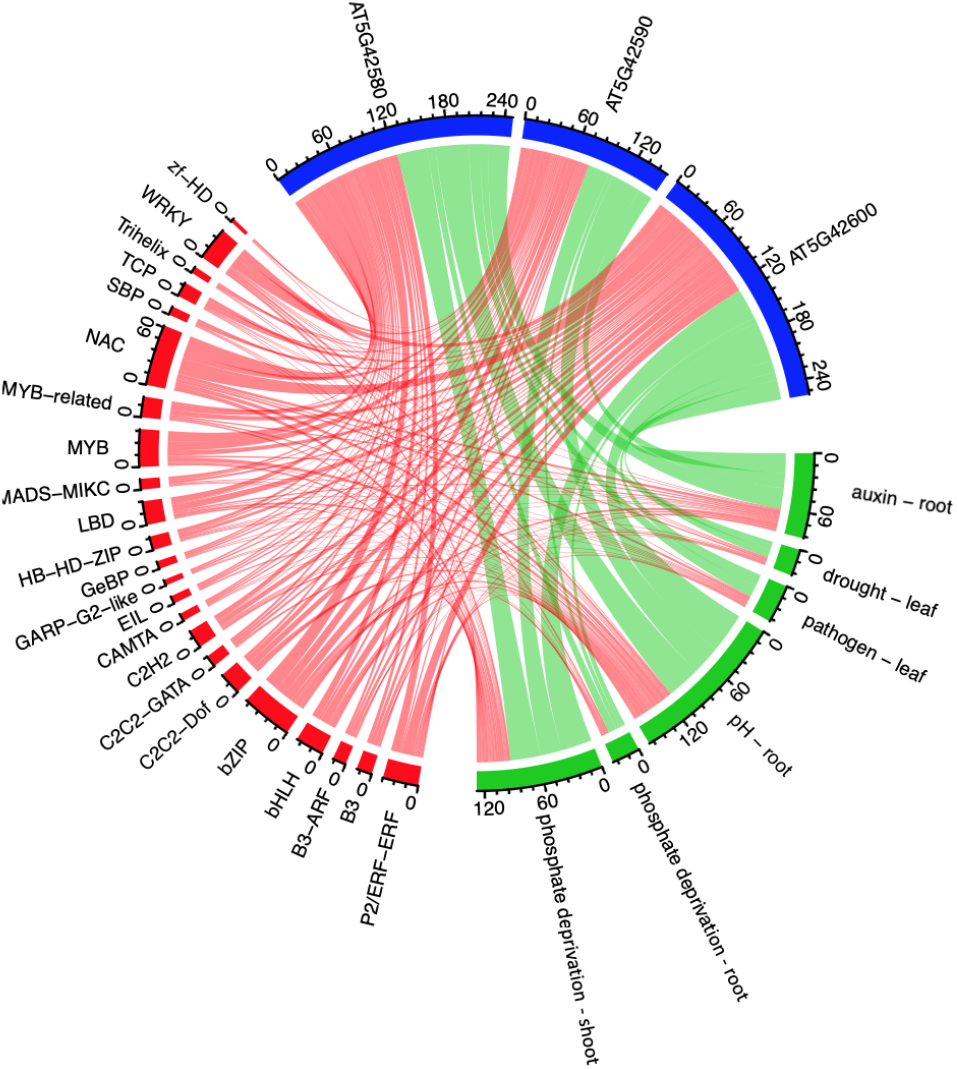
Cluster gene level visualization, i.e. transcription factor families (red) vs conditions and treatments (green) vs cluster genes (blue), of transcriptional activity and regulation of the marneral gene cluster.

**Figure 4:**
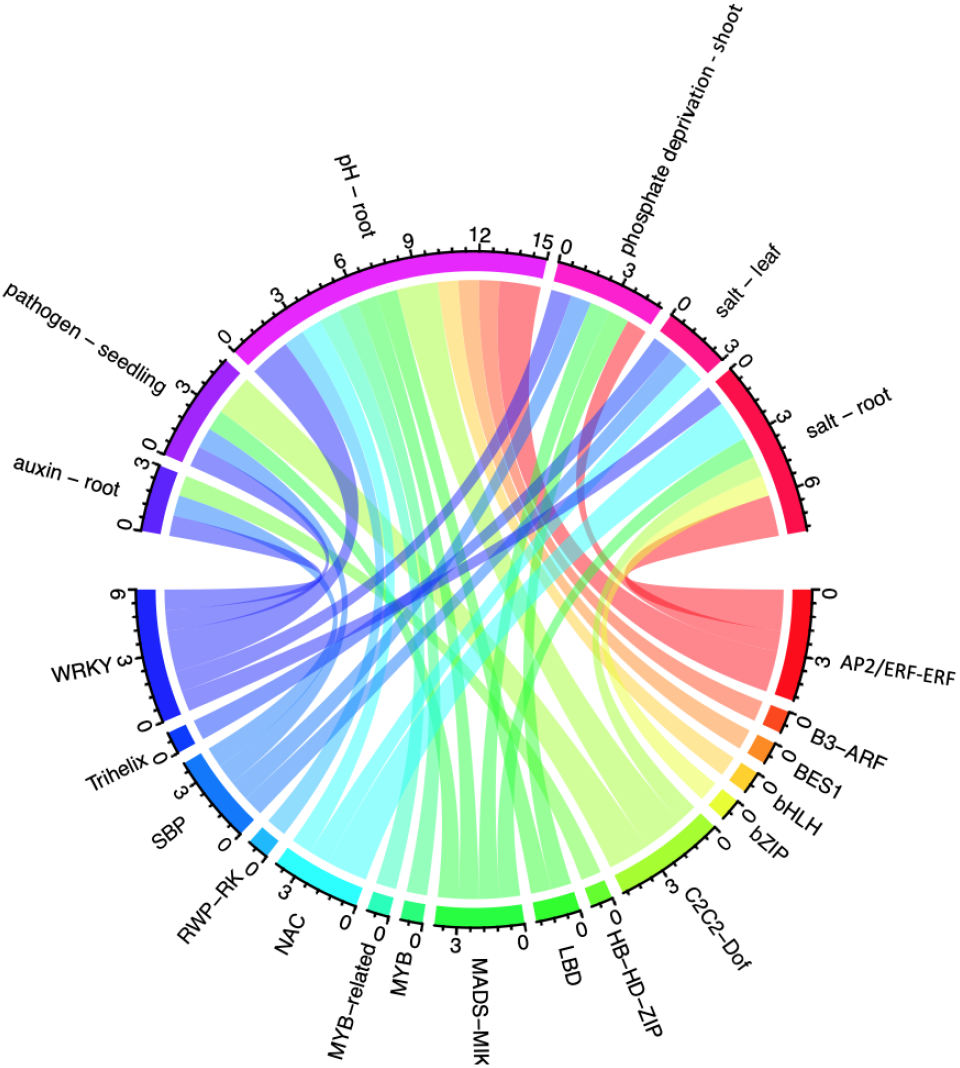
Gene cluster level visualization, i.e. transcription factor families vs conditions and treatments, of transcriptional activity and regulation of the thalianol gene cluster.

**Figure 5:**
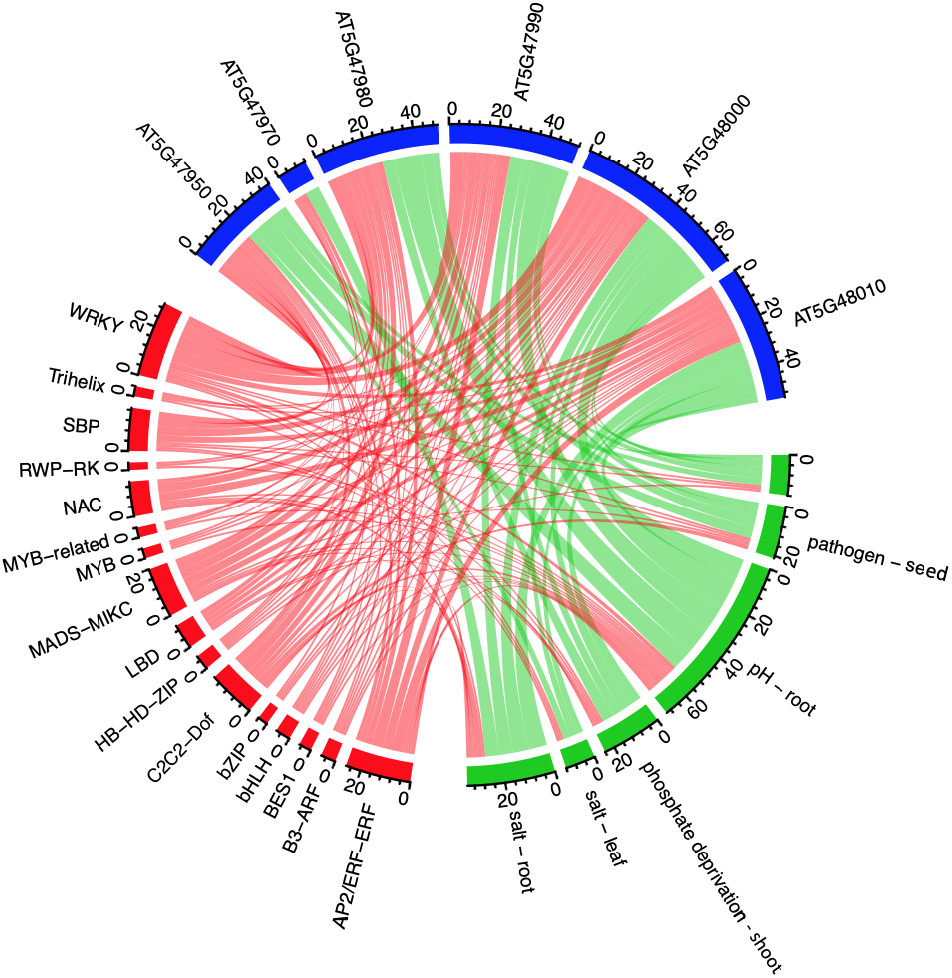
Cluster gene level visualization, i.e. transcription factor families (red) vs conditions and treatments (green) vs cluster genes (blue), of transcriptional activity and regulation of the thalianol gene cluster.

## 3. Results and Discussion

To demonstrate the utility of METACLUSTER^*plus*^ as well as its visualization capabilities, we run our pipeline for metabolic gene cluster predictions in *Arabidopsis thaliana*, acquired from [13]. Here, we highlight prediction and visualization of transcriptional activity and regulation for two examples, the experimentally characterized terpene biosynthetic clusters in Arabidopsis, i.e., the thalianol [7] and the marneral [8] cluster. These were clusters C641 and C628 in [13]. We use a recently compiled large-scale gene expression dataset by He *et al*. [9] with 6057 expression profiles, covering 79.7% of the *A. thaliana* ecotype Columbia genome. We retain 435 experimental treatments represented by 1825 expression profiles measuring gene expression responses of wild type plants to treatment and control conditions. All 435 experimental treatments are assigned to 27 manually curated conditions and 9 tissues (see [1], supplementary methods). As for a gene regulatory network architecture, we harness a recently released, large scale DNA affinity purification sequencing (DAP-seq) based dataset [2], providing a gene regulatory network consists of 349 transcription factors, 26921 target genes and 1791998 connections.

Table 1 illustrates an excerpt of the textual representation of the predicted gene cluster regulations. Figures 2 and 4 as well as figures 3 and 5 highlight the corresponding chord graph based visualizations on a gene cluster as well as cluster specific gene levels, respectively. In particular, these graphical representations may serve to provide an immediate visual and high level summary of the given condition-specific regulatory relationships for prioritization and further investigations. For instance, both clusters show an enrichment for regulation by members of the APETALA2/Ethylene Response Factor (AP2/ERF) family, or the WRKY transcription factor family across a variety of stress conditions and tissues, which is corroborated by research on these transcription factor families’ influence on specialized metabolism control in plants [20, 19, 18].

Given its utility, we anticipate METACLUSTER^*plus*^ to be a valuable tool for the efficient integration of heterogeneous datasets in order plan experiments and guide validation of context-specific metabolic gene cluster transcriptional regulation.

## Notes

### Competing Interest Statement

The authors have declared no competing interest.

